# Crystal structure of *Sulfolobus solfataricus* topoisomerase III reveals a novel carboxyl-terminal zinc finger domain essential for decatenation activity

**DOI:** 10.1101/706986

**Authors:** Hanqian Wang, Junhua Zhang, Zengqiang Gao, Xin Zheng, Keli Zhu, Zhenfeng Zhang, Zhiyong Zhang, Yuhui Dong, Li Huang, Yong Gong

## Abstract

DNA topoisomerases are essential enzymes for a variety of cellular processes involved in DNA transactions. Mechanistic insights into type IA DNA topoisomerases have come principally from studies on bacterial and eukaryotic enzymes. A structural understanding of type IA topoisomerases in Archaea is lacking. Here, we present a 2.1-angstrom crystal structure of full-length *Sulfolobus solfataricus* topoisomerase III (*Sso* topo III), an archaeal member of type IA topoisomerases. The structure shows that *Sso* topo III adopts a characteristic torus-like architecture consisting of a four-domain core region and a novel carboxyl-terminal zinc finger domain (domain V). Structure-based mutation analyses reveal that a novel zinc-binding motif in domain V is essential for the DNA decatenation activity of *Sso* topo III. Our data indicate that *Sso* topo III represents a subclass of Type IA topoisomerases capable of resolving DNA catenates using a domain V-dependent mechanism.

**IMPORTANCE:** Type IA topoisomerases are omnipresent in all cellular life forms and serve pivotal roles in cellular processes involved in DNA transactions. While considerable insights have been gained into Type IA topoisomerases from bacteria and eukaryotes, a structural understanding of type IA topoisomerases in Archaea remains elusive. we first determined the crystal structure of full-length *Sulfolobus solfataricus* topoisomerase III (*Sso* topo III), an archaeal member of type IA topoisomerases. Our structure provides the first molecular view of this archaeal topoisomerase, which removes negative supercoils and decatenates DNA catenane. Our findings manifest that *Sso* topo III may serve as an alternative prototype of type IA topoisomerases, whose decatenation mechanism differs from that of known bacterial and eukaryotic topoisomerases III such as *Escherichia coli* topoisomerase III (EcTOP3).

## INTRODUCTION

The overall topological state of DNA in cells is altered by DNA topoisomerases. In all three domains of life, conserved DNA topoisomerases are essential for a wide variety of cellular processes involved in DNA transactions (1–4). There are two types of topoisomerases (1). Type I topoisomerases cut only one strand of DNA, while type II topoisomerases cleave both strands of the DNA duplex. Type I enzymes can be further classified into three different families, types IA, IB and IC, on the basis of transiently formed covalent intermediates formed (5′- or 3′- covalent intermediate) (5). *Escherichia coli* topoisomerase I (EcTOP1) and topoisomerase III (EcTOP3) are the best-characterized bacterial type IA enzymes to date.

A conserved toroidal fold formed by four domains (I-IV) has been found in the structures of type IA enzymes including EcTOP1 (6), EcTOP3 (7), *Thermotoga maritima* topoisomerase I (TmTOP1) (8), *Mycobacterium tuberculosis* topoisomerase I (MtTOP1) (9), *Mycobacterium smegmatis* topoisomerase I (MsTOP1) (10), human topoisomerase IIIα (HsTop3α) (11), human topoisomerase IIIβ (HsTop3β) (12) and a reverse gyrase (13) from *Archaeoglobus fulgidus*, a hyperthermophilic archaeon. Structural studies (14–16) as well as biochemical (17–20) and single-molecule (21–24) experimental data have provided abundant evidences in support of an enzyme-bridged strand passage model for type IA enzymes. According to this model, a type IA topoisomerase nicks one DNA single-strand, and remains attached to the two ends of the nick. That is, the enzyme is not only covalently attached to one end of the nick, but also noncovalently bound to the other end to generate a bridge through which another single-strand or the DNA duplex is passed. The following step is the resealing of the nick and the release of the enzyme. In this process, the OH group of the aromatic ring of the Tyr residue in the enzyme nucleophilically attacks the scissile phosphodiester bond, producing a transient 5′ phosphotyrosine covalent bond.

Members of the type IA subfamily are further divided into two mechanistically distinct subclasses based on their biochemical specificity for substrates (2). EcTOP1 and EcTOP3 are the prototypes of these two subclasses of type IA enzymes. EcTOP3 catalyzes the DNA catenates more effectively than it relaxes negatively supercoiled DNA, whereas the converse is true for EcTOP1 (23, 25–28). EcTOP3, but not EcTOP1, contains a unique loop located close to the central hole (29). This positively charged loop, known as the decatenation loop, is essential for the decatenation activity of EcTOP3 (24, 29).

Previous studies have characterized topoisomerase III (hereafter, *Sso* topo III) from *Sulfolobus solfataricus*, a hyperthermophilic archaeon, as a member of the type IA topoisomerase family (30, 31). A phylogenetic analysis of *Sso* topo III, and the observation that *Sso* topo III inefficiently catalyzes the relaxation of negatively supercoiled DNA, led to the suggestion that *Sso* topo III is more similar to EcTOP3 than to EcTOP1 (30). More recently, Bizard et al. provided convincing biochemical data in support of notion that *Sso* topo III is a genuine decatenase that can efficiently unlink covalently closed catenanes alone at temperatures up to 90°C (32). However, little is known about the structure of type IA topoisomerases (except for reverse gyrase) from the Archaea. The structural basis for the decatenase activity of *Sso* topo III is unclear.

Here, we have determined the crystal structure of full-length *Sso* topo III. *Sso* topo III adopts a classical elongated toroidal-like shape with four domains (I-IV) forming the protein core and a novel carboxyl-terminal zinc finger domain (domain V) tethered to the core against domain II. Furthermore, domain V is conserved among DNA topoisomerases III from the TACK superphylum of the Archaea. In addition, structure-based mutational analyses show that, whereas the zinc-binding motif of domain V was required for efficient DNA relaxation by *Sso* topo III, its depletion nearly abolished the DNA decatenation activity of the enzyme. These findings indicate that *Sso* topo III may represent as a novel type of topoisomerases III which employ a decatenation mechanism distinctly different from that of known bacterial and eukaryotic equivalent.

## RESULTS

### Overall structure of intact *Sso* topo III

To obtain crystals of *Sso* topo III, we overproduced and purified sufficient quantities of the mutant enzyme (Y318F) in which the catalytic residue Tyr318 was replaced with Phe (Fig. 1 A). The structure of the full-length enzyme was determined at 2.1 Å resolution using molecular replacement (Table 1). The full-length protein formed an asymmetric unit comprising four *Sso* topo III molecules. The four molecules had essentially the same architecture. For convenience, one molecule was chosen for the following structural analyses.

**FIG 1.**
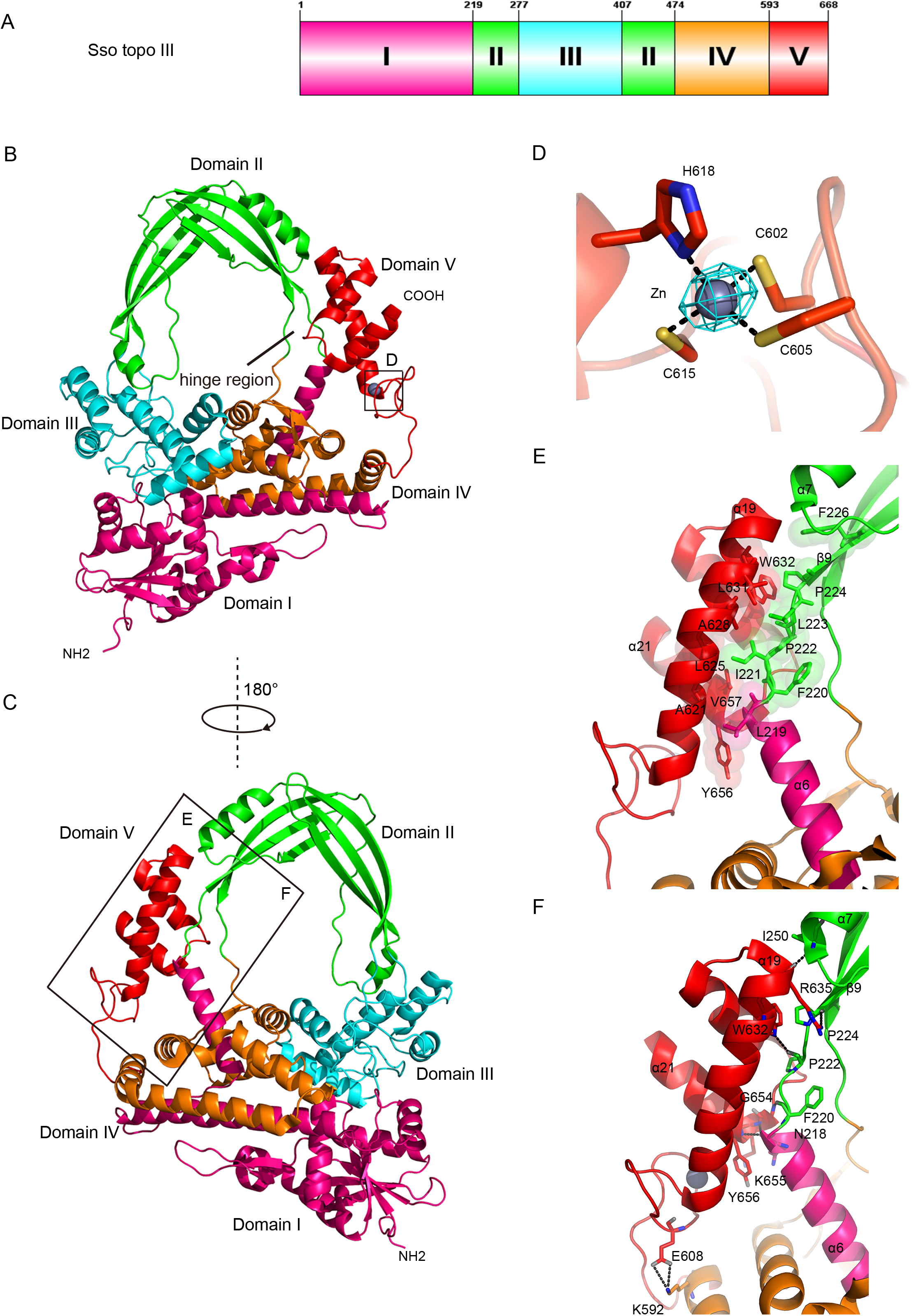
Overall structure of full-length *Sso* topo III in apo state. A. Domain organization of *Sso* topo III. *Sso* topo III is composed of N-terminal four domains (domain I-IV) and the unique carboxyl-terminal domain V. Domains I-V are colored hotpink, green, cyan, orange, red, respectively. B. C. Two views of *Sso* topo III. Domains I-V are colored as the legend to Figure 1A. The close-up view (D) is highlighted by the solid box in B. Detailed interactions (E and F) within the domains II/V interface is highlighted by the solid box in C. D. The close-up view shows the anomalous difference electron density map (drawn in cyan mesh) around Zn (II) contoured at 15σ using the diffraction data collected at the Zn absorption peak. The three Cys (C602, C605, C615) and one His residue (H618) coordinating Zn (II) are shown in sticks. Zn (II) is shown as a gray sphere. E. Hydrophobic zipper between domain V residues from helix α19 and domain II residues from the loop linking domain I and II. The hydrophobic interactions are shown in space-filling presentations. F. As in E but describing the corresponding hydrogen-bonding network. The hydrogen bonds are shown as black dashed lines.

**Table 1.**
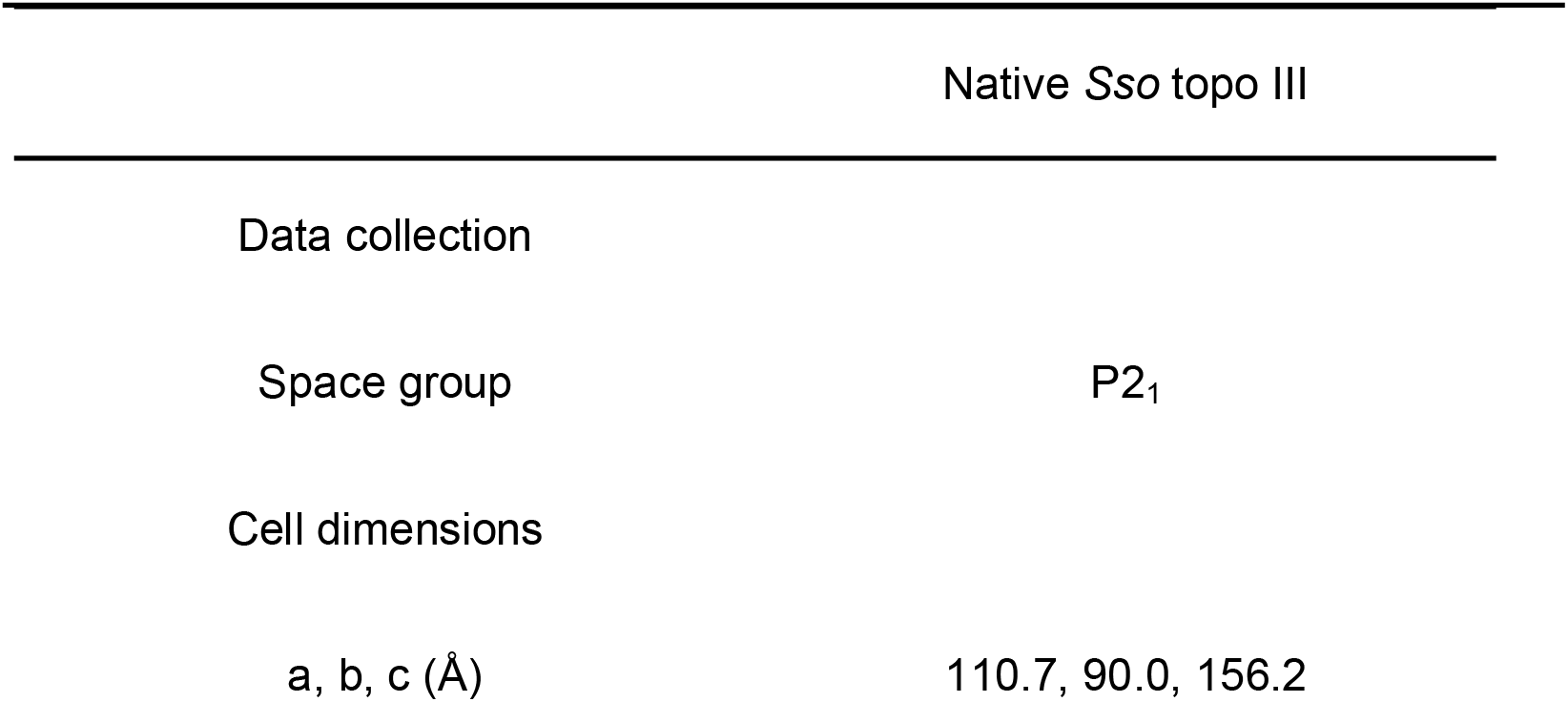
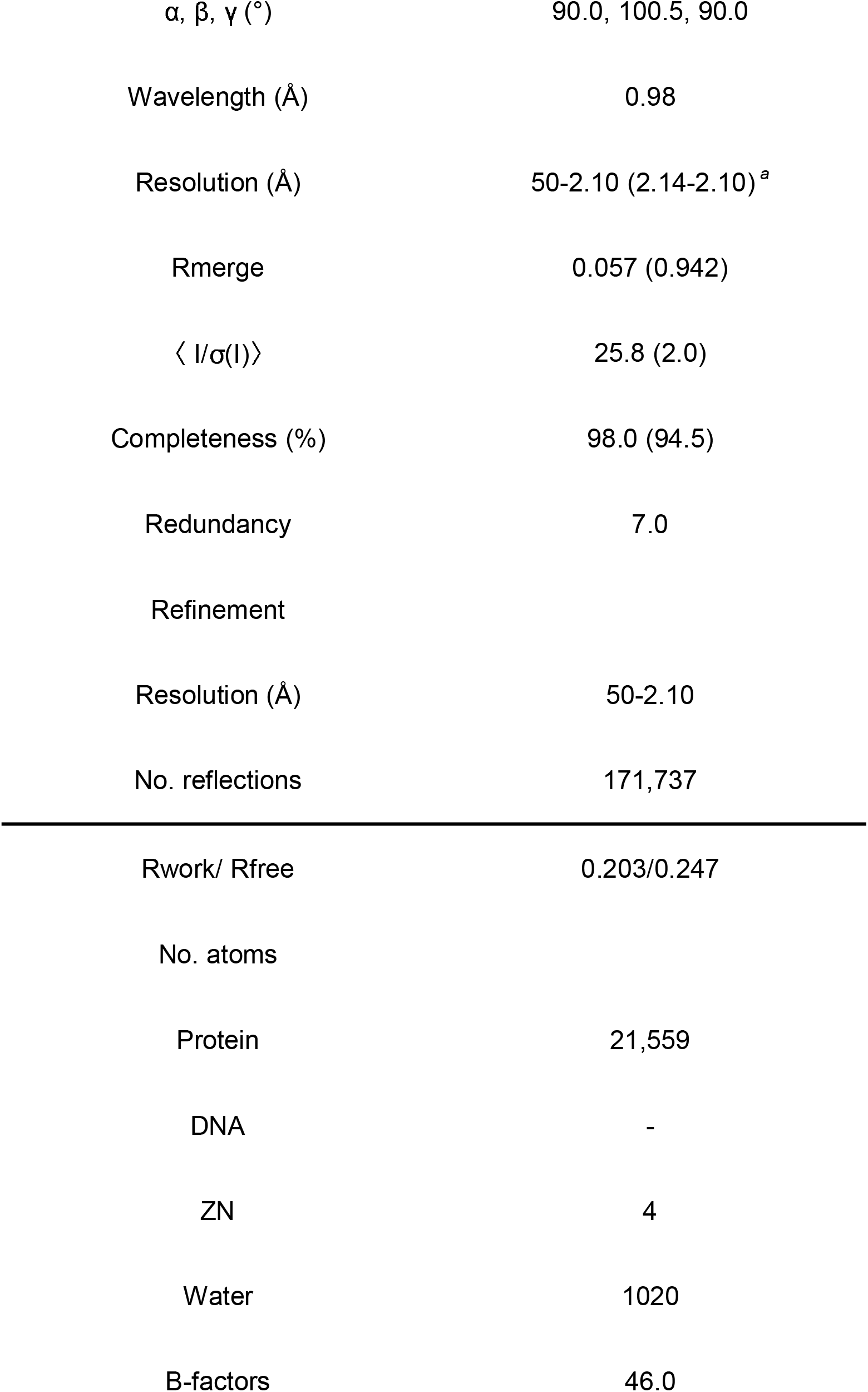
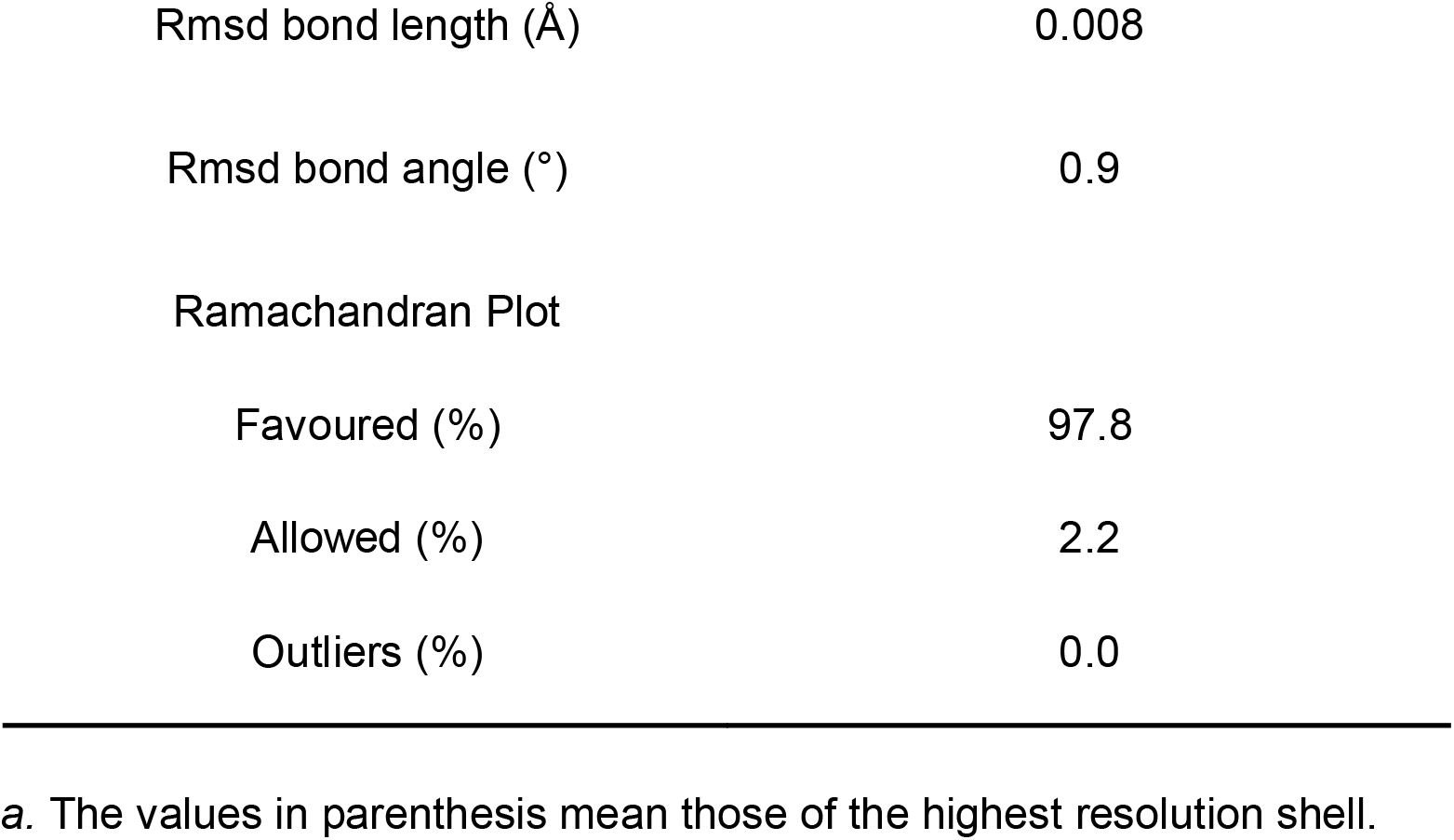
Data collection and refinement statistics

Our analysis showed that the overall structure of *Sso* topo III comprises four domains (I–IV) (residues 1–593), forming the conserved core, and a novel carboxyl-terminal domain V (residues 594–668) (Fig. 1A, 1B and S1). The enzyme adopts a canonical toroidal-like elongated configuration with a large hole (Fig. 1B and 1C), which is shared by all known type 1A topoisomerases (33). The enzyme-bridged strand passage model proposes that the hole accommodates single or double-stranded DNA (34).

Domain I (residues 1–219) exhibits a topoisomerase–primase fold (2), which is found in type IA, IIA, and IIB topoisomerases, as well as in bacterial primases. Within domain I, a disulphide bridge crosslinks Cys4 of a loop (residues 1–10) and Cys34 of an α/β fold consisting of six parallel β strands sandwiched with four α helices (Fig. S1). This connection results in tight packing of domain I and stabilizes its architecture. In Domain II (residues 220–277, 408–474), the hinge region (residues 218–224, 471–480) is composed of two loops linking domains I and II, and II and IV, respectively (Fig. 1B). The hinge is believed to serve as the principal pivot point for opening and closing of the DNA gate in the strand passage reaction (6, 11, 35).

A long loop (residues 594–616) leads from helix α18 of domain IV to domain V (residues 594–668), thus linking the core of the enzyme to domain V (Fig. 1B and S1). Domain V attaches to the top of the topoisomerase (domain II) against the hinge region (Fig. 1B and 1C). Three antiparallel α-helices, including helix α19 (residues 616–636), helix α20 (residues 640–649), and helix α21 (residues 655–666), are coiled around each other to constitute the bulk of domain V (Fig. 1B). Interestingly, four residues (Cys602, Cys605, Cys615 and His618) coordinate with a zinc atom to generate a novel zinc-binding motif (Fig. 1D and S1). Among them, the three Cys residues come from the long loop connecting domains IV/V and the His residue is from helix α19. The presence of the znic-binding site was confirmed by an analysis of the anomalous difference Fourier map (Fig. 1D). The recently characterized mesophilic archaeal topoisomerase, *Methanosarcina acetivorans* topo IIIα (*Mac* topo IIIα), also has one znic-binding site (36). However, the zinc-binding motif of *Sso* topo III is the Cys3His1 type, whereas that of *Mac* topo IIIα is tetraCys type. For *Sso* topo III, the zinc finger motif may contribute to the stabilization of domain V. As revealed by sequence alignment, the carboxyl-terminal zinc finger motif of the Cys3His1 type is highly conserved among topoisomerases from the Crenarchaeota (Fig. S2A), suggesting that this motif plays an important role. Intriguingly, small proteins homologous to domain V are also found in Thaumarchaeota and Bathyarchaeota, both of which are phyla within the TACK superphylum where Crenarchaeota also belong, as well as Euryarchaeota and Thorarchaeota (Fig. S3). Homologues of *Sso* topo III are present in Thaumarchaeota, Bathyarchaeota, Euryarchaeota and Thorarchaeota, but these enzymes almost lack the carboxyl-terminal domain similar to domain V in *Sso* topo III. It would be of interest to determine if the domain V-like small proteins physically and functionally interact with the type IA topoisomerases in these organisms. Notably, no structural elements similar to domain V of *Sso* topo III is found in the Protein Data Bank, as detected by DALI (37). These results suggest that the carboxyl-terminal fold of *Sso* topo III, also found in several archaeal lineages, is unique.

### Interface of domains II and V

A distinguishing feature of the overall structure of *Sso* topo III compared with that of other known available type IA topoisomerases is the presence of three helices in domain V located close to the hinge region of domain II (Fig. 1B). Two flexible loops, each about five residues long, link these three helices. This arrangement causes the three helices to pack against each other so that their movement is highly coordinated, but they maintain a degree of flexibility. Additionally, within the long linker loop (about 20 residues) connecting domains IV and V, there are three successive turns, which maintain free space between the two domains. The novel zinc-binding motif, located on the carboxyl-end of the long loop, contributes to the stabilization of the loop (Fig. 1D). By analogy to the other solved type IA topoisomerases (6), the interfaces between domain III and domains IV and I in *Sso* topo III are conserved and far-ranging. However, the interface between domains II and V within the enzyme is unique.

The longest one of the three helices in domain V, helix α19, acts as the major binding platform for the interface between domains II and V. The interface is mostly, but not exclusively, hydrophobic (Fig. 1E). Extensive hydrophobic interactions mainly occur between the side chains of Leu219, Phe220, Ile221, Pro222, Leu223, Pro224 and Phe226 in the hinge region, which connects helix α6 of domain I with strand β9 of domain II, and the side chains of Ala621, Leu625, Ala628, Leu631, Trp632 in helix α19 and Tyr656, Val657 in helix α21 of domain V (Fig. 1E). Moreover, the main chain amides of Gly654 and Lys655 in domain V form hydrophilic interactions with the backbone carbonyl groups of Phe220 and Asn218 in the hinge region, respectively, while the amide of Tyr656 makes hydrophilic interaction with the backbone carbonyl groups of Asn218 (Fig. 1F). In addition, the NH group on an indole group of Trp632 makes a hydrogen bond with the backbone carbonyl group of Pro222. These interactions further stabilize the attachment of domain V to domain II. Most importantly, the main chain carbonyl group and side chain amide of Arg635 in the carboxyl-end of helix α19 in domain V form hydrogen bonds with the main chain amide of Ile250 at the amino-end of helix α7 and the carbonyl group of Pro224 at the amino-end of strand β9 in domain II, respectively (Fig. 1F). Domain V and the main part of domain II are connected by a single interaction, that is, double hydrogen bonds involving Arg635. This may allow for independent and coordinated movements of domain V relative to domain II at different stages during catalysis. Noticeably, Glu608 from domain V forms salt bridge with Lys592 from domain IV on the other end of the interface. Alignment of the sequences of type IA topoisomerases from the Crenarchaeota shows that most of the above-mentioned residues contributing to the interaction between domains II and V are highly conserved (Fig. S2B). This implies that the interaction between domains II and V may be essential for the functional role of domain V of the archaeal topoisomerases.

### Comparison of the interactions around the hinge region among type IA topoisomerases

The studies on EcTOP3 (7, 14, 29) indicated that the decatenation loop on domain IV is critical for efficient DNA decatenation (Fig. 2A). A recent report on *Sso* topo III provides strong biochemical evidence that the enzyme is a true decatenase (32). Surprisingly, the structural equivalent of the decatenation loop is absent from *Sso* topo III (Fig. 2A), suggesting that DNA decatenation by *Sso* topo III may differ mechanistically from that by EcTOP3.

**FIG 2.**
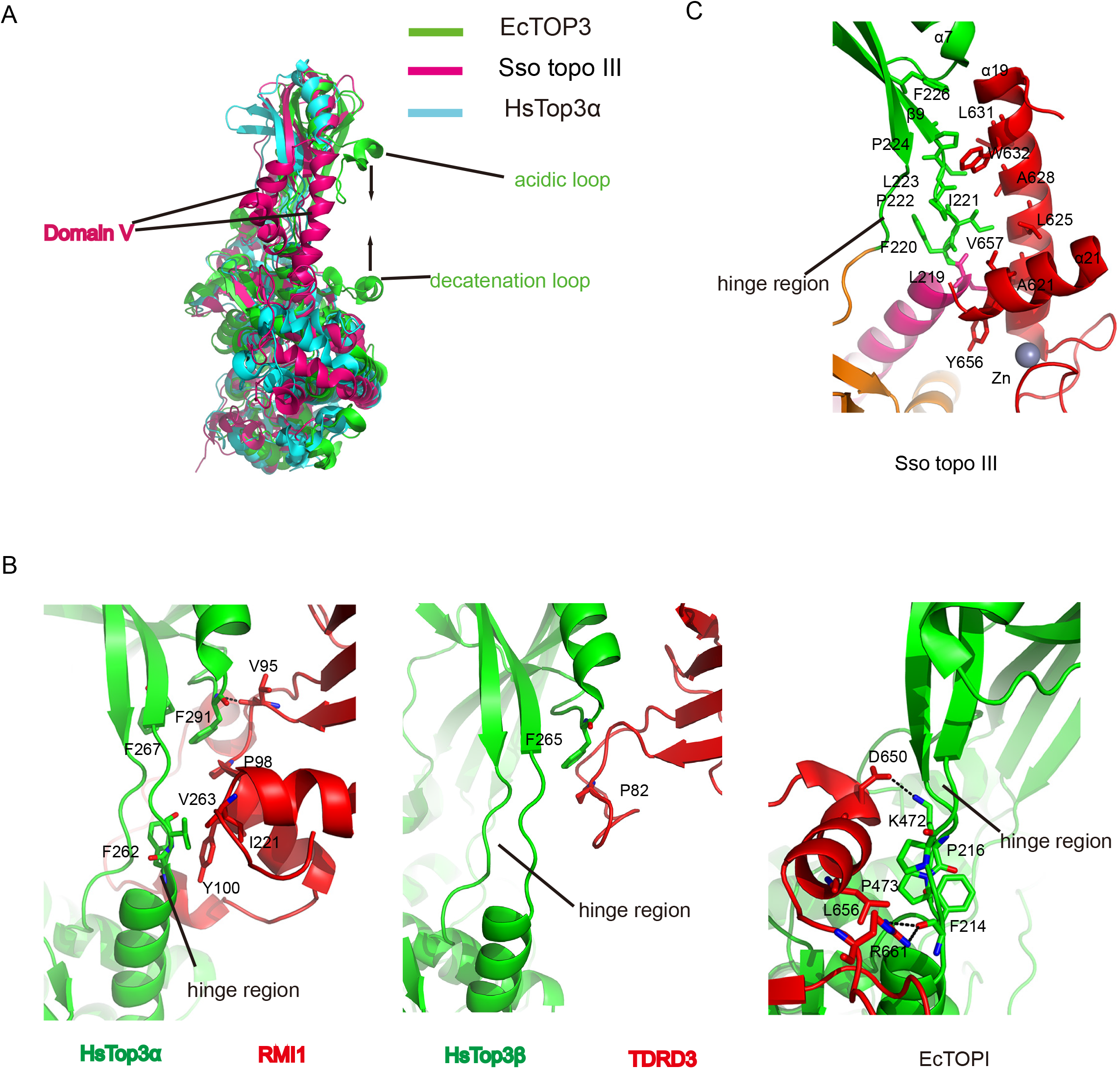
Comparison of structural elements around the hinge region in *Sso* topo III, EcTOP1, EcTOP3, HsTop3α and HsTop3β. A. Overall structural comparison among *Sso* topo III (hotpink), EcTOP3 (green) and HsTop3α (cyan). For clarity, the RMI1 subunit of HsTop3α complex (PDB entry 4CHT) is not shown. Domain V of *Sso* topo III is indicated. The decatenation loop and the acidic loop of EcTOP3 (PDB entry 1D6M) are shown, and their movement during the gate dynamics is shown with opposing arrows. B. The close-up views around the hinge regions of HsTop3α in Top3α-RMI1 complex (PDB entry 4CHT), HsTop3β in Top3β-TDRD3 complex (PDB entry 5GVE) complexes, and EcTOP1, mainly showing the hydrophobic interactions. The hydrogen bonds are shown as black dashed lines. For clarity, the ssDNA molecule of EcTOP1 complex (PDB entry 4RUL) is not shown. C. The close-up view around the hinge region of *Sso* topo III, showing the hydrophobic interactions. For clarity, helix α20 (residues 639-654) of *Sso* topo III is not shown.

It is believed that the highly conserved hinge region joining domains II and IV appears to be important for strand passage during catalysis (6). To understand the decatenation mechanism of *Sso* topo III, we performed a structural analysis of the potential interactions within the available type IA topoisomerases between the hinge region and the carboxyl-terminal domain or an additional subunit of the topoisomerase complex. No direct interactions exist between the hinge regions and the C-terminus in TmTOP1 (8), MtTOP1 (9) and MsTOP1 (10). However, there are direct interactions between the hinge region and parts of the carboxyl-terminal domain in several topoisomerases or the other subunit of a topoisomerase complex (Fig. 2B). The interactions may influence the opening state of their gates. The human BLM helicase, topoisomerase IIIα (HsTop3α) and RMI1 proteins form a minimal protein complex, which catalyses the dissolution of double-Holliday junctions (dHJ) to produce non-crossover products (38). A structural analysis of the Top3a–RMI1 complex showed that the hinge region makes intimate contacts with the decatenation loop of the RMI1 subunit by hydrophobic interaction, suggesting that this contact contributes to the modulation of the opening-closing state of the gate in Top3α (11) (Fig. 2B). Similar to the Top3a–RMI1 complex, Top3β forms a complex with the TDRD3 subunit. A structural analysis of the complex indicated that the hinge region of Top3β is a little bit apart from the insertion loop of TDRD3 (12) (Fig. 2B). In the case of EcTOP1, the interaction between a unique α helix in domain VI of the carboxyl-terminal domain and the hinge region mediates the major direct contact between the carboxyl-terminal domain and the catalytic core (39) (Fig. 2B). The conformational changes of the carboxyl-terminal domain are potentially transferred to the hinge region through the unique helix in domain VI at some points in the catalytic cycle, thereby regulating the opening-closing state of the gate. In *Sso* topo III, major contacts between the hinge region and the longer helix α19 of domain V occur through stable and extensive hydrophobic interactions (Fig. 2C), and several hydrogen bonds, located at the two ends of the interface between domains II and V, are involved in the interaction between domain V and the bulk of domain II and domain IV, respectively (Fig. 1F). This network of interactions may allow domain V to adhere to the hinge region mainly by the α19 helix during the strand passage reaction, and possibly make the bulk of domain II and domain IV to move relative to the hinge region at some points during the catalysis.

### Deletion of the zinc-binding motif reduces the relaxation activity of *Sso* topo III

To explore the role of the carboxyl-terminal domain, the zinc-binding motif in particular, in DNA relaxation by *Sso* topo III, a carboxyl-terminally truncated mutant (597ΔC), which lacked whole domain V, and a quadruple mutant (CCCH/4A), in which the four residues (Cys602, Cys605, Cys615 and His618) coordinating with the zinc atom were all replaced with Ala, were generated by mutagenesis (Table S1). In the CCCH/4A, the mutation of the zinc-binding motif (3Cys1His type) resulted in the elimination of the zinc atom (Fig. 1D). These two mutant proteins were overproduced in *E. coli* and purified to homogeneity. The two mutant proteins showed the same optimal temperature as the wild-type enzyme (75°C) in DNA relaxation (Fig. S4). 597ΔC possessed about 50% of the wild-type activity (Fig. 3, compare top and middle panel), whereas CCCH/4A retained about 15% of the wild-type activity (Fig. 3, compare top and bottom panel). These results indicated that the zinc-binding motif is necessary for efficient relaxation by *Sso* topo III.

**FIG 3.**
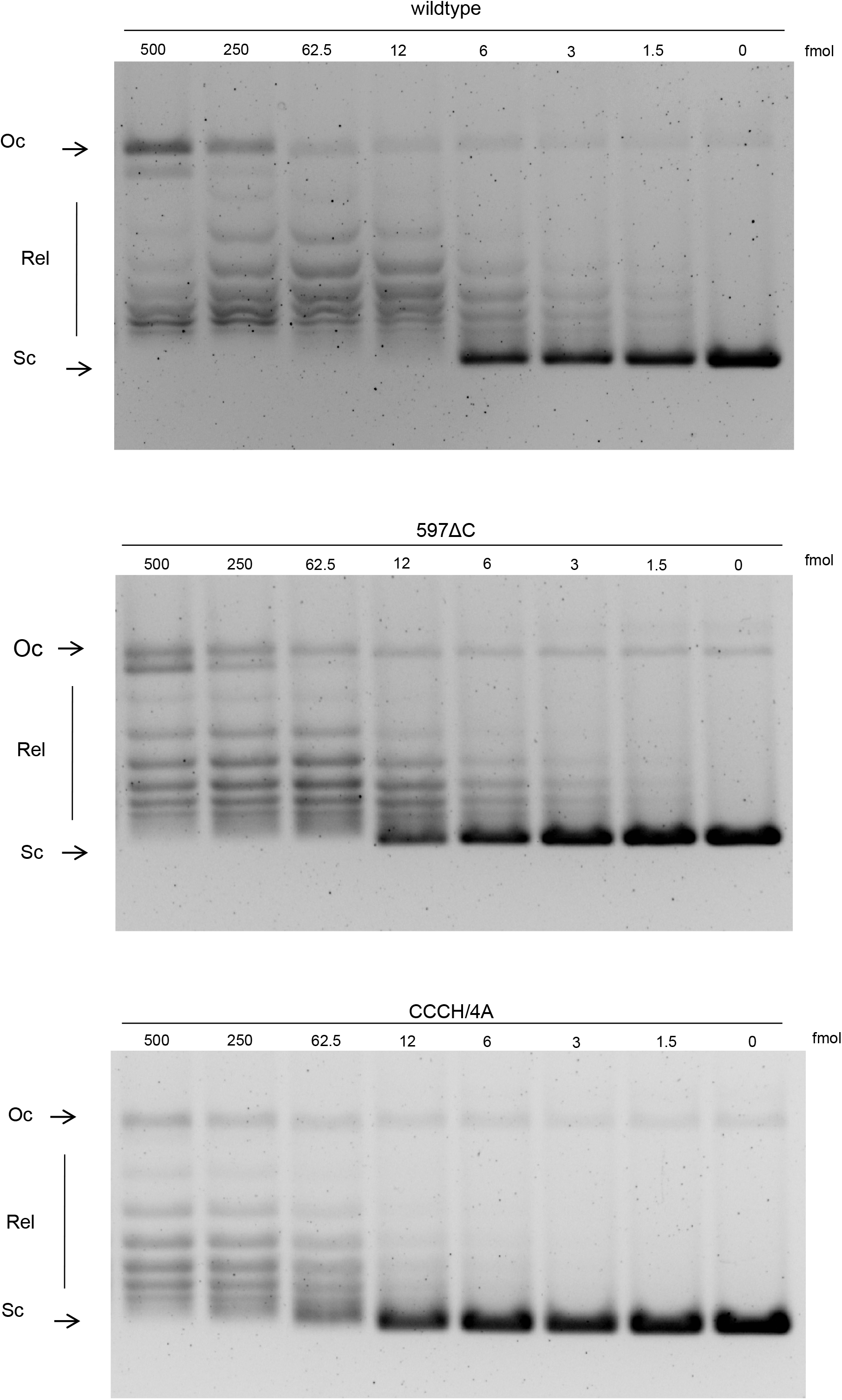
Relaxation activity of wild-type and mutant *Sso* topo III proteins. pUC18 DNA was incubated at 75 °C for 30 min in relaxation reaction buffer (50 mM Tris-HCl (pH 8.8), 20 mM MgCl2, 1 mM DTT, 0.1 mM EDTA, 90 mM sodium chloride, 30 μgml^-1^ BSA, and 12% (vol/vol) ethylene glycol)) with wild-type *Sso* topo III (top panel), 597ΔC (middle panel) and CCCH/4A (bottom panel) at 500, 250, 62.5, 12, 6, 3, 1.5 and o fmol. Reactions were terminated; the DNA products were separated in 1.4% agarose gels and visualized by staining with ethidium bromide. Oc indicates open circle nicked or gapped circular DNA; Sc denotes negatively supercoiled circular DNA; and Rel is the relaxed topoisomers.

To compare DNA relaxation reactions catalysed by wild-type and mutants of *Sso* topo III, a time-course assay was performed (Fig. 4). The wild-type protein displayed a profile of topoisomers indicative of a distribute mechanism of relaxation (Fig. 4, top panel), as described recently (32). However, both 597ΔC and CCCH/4A exhibit a relaxation profile differently from the wild-type *Sso* topo III (Fig. 4, compare top, middle, and bottom panel). The DNA relaxation reaction catalysed by 597ΔC appeared to more distributive than that by the wild-type protein. Interestingly, the relaxation reaction becomes more distributive, when the unique zinc-binding motif was eliminated, than that as the carboxyl-terminal domain of *Sso* topo III was wholly deleted (Fig. 4). Together, these results indicated that domain V, especially the zinc-binding motif, facilitates the relaxation of negatively supercoiled DNA by *Sso* topo III.

**FIG 4.**
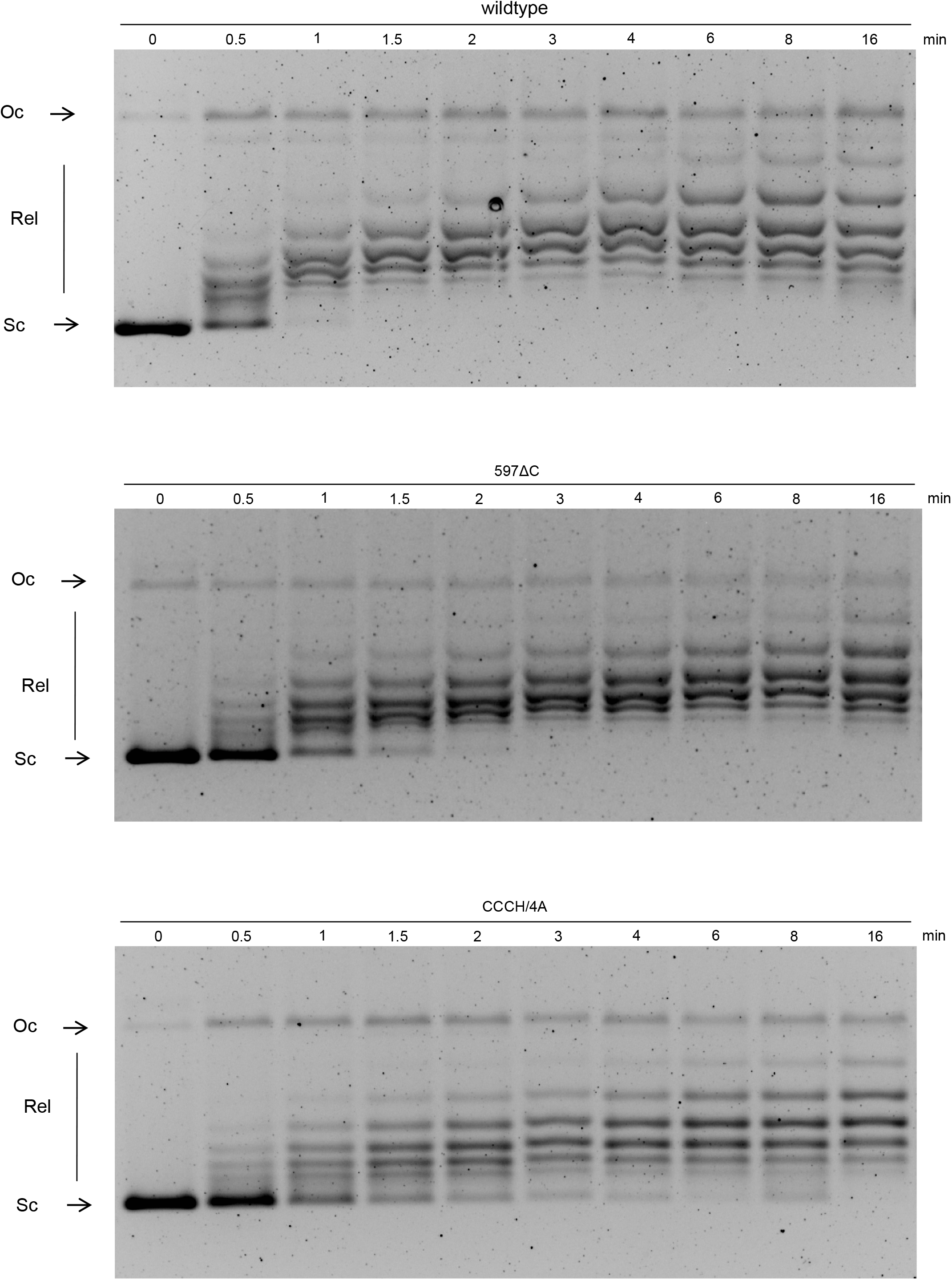
Time-course of DNA relaxation by wild-type and mutants *Sso* topo III proteins. DNA relaxation by wild-type *Sso* topo III (top panel), 597ΔC (middle panel) and CCCH/4A mutant (bottom panel), were carried out over the indicated time periods as described in the FIG 3. The molar ratio of protein to pUC18 DNA was 1:1 in the reaction. Samples terminated at the indicated time-points. Sample at o-min point indicated untreated substrate pUC18 DNA. The reaction products were analyzed in 1.4% agarose gels and visualized by staining with ethidium bromide. Oc indicates open circle nicked or gapped circular DNA; Sc denotes negatively supercoiled circular DNA; and Rel is the relaxed topoisomers.

To probe the potential roles of domain V in *Sso*Topo III, we also determined ssDNA binding by the domain V-deletion mutant protein 594ΔC (residues 1-594). The mutant protein showed a ~ 5-fold lower affinity than wild-type *Sso*Topo III for a 32-nt ssDNA containing the preferred cleavage site (designated C32) than the wild-type protein (Fig. S5). Therefore, it appeared that domain V contributes significantly to template binding by *Sso*Topo III.

### The novel zinc-binding motif is essential for DNA decatenation by *Sso* topo III

To explore the role of the novel domain V in *Sso* topo III-catalysed decatenation, wild-type and mutants of *Sso* topo III were subjected to a decatenation assay using kinetoplast DNA (kDNA) (40) as the substrate. Wild-type *Sso* topo III was optimally active in kDNA decatenation at 90°C, in consistent with the observation made previously (32) (Fig. S6A, B, C). The fast-migrating bands, corresponding to closed minicircles, were present when the reactions were performed at very high temperatures (90 °C) (Fig. 5A). These closed minicircles were released from the kDNA network by *Sso* topo III, because the enzyme apparently was unable to relax supercoiled DNA at 90 °C and above, but was still able to decatenate DNA catenanes (32). Therefore, the presence of closed minicircles at very high temperature was considered as a measure of the decatenation activity of *Sso* topo III. The bands corresponding to closed minicircles were undetectable in reactions containing 597ΔC (Fig. 5B), suggesting that the mutant protein had no detectable decatenation activity at very high temperature. Similarly, CCCH/4A was nearly inactive in decatenation (Fig. 5C). Taken together, we concluded that domain V, the zinc-binding motif in particular, is essential for the decatenation activity of *Sso* topo III.

**FIG 5.**
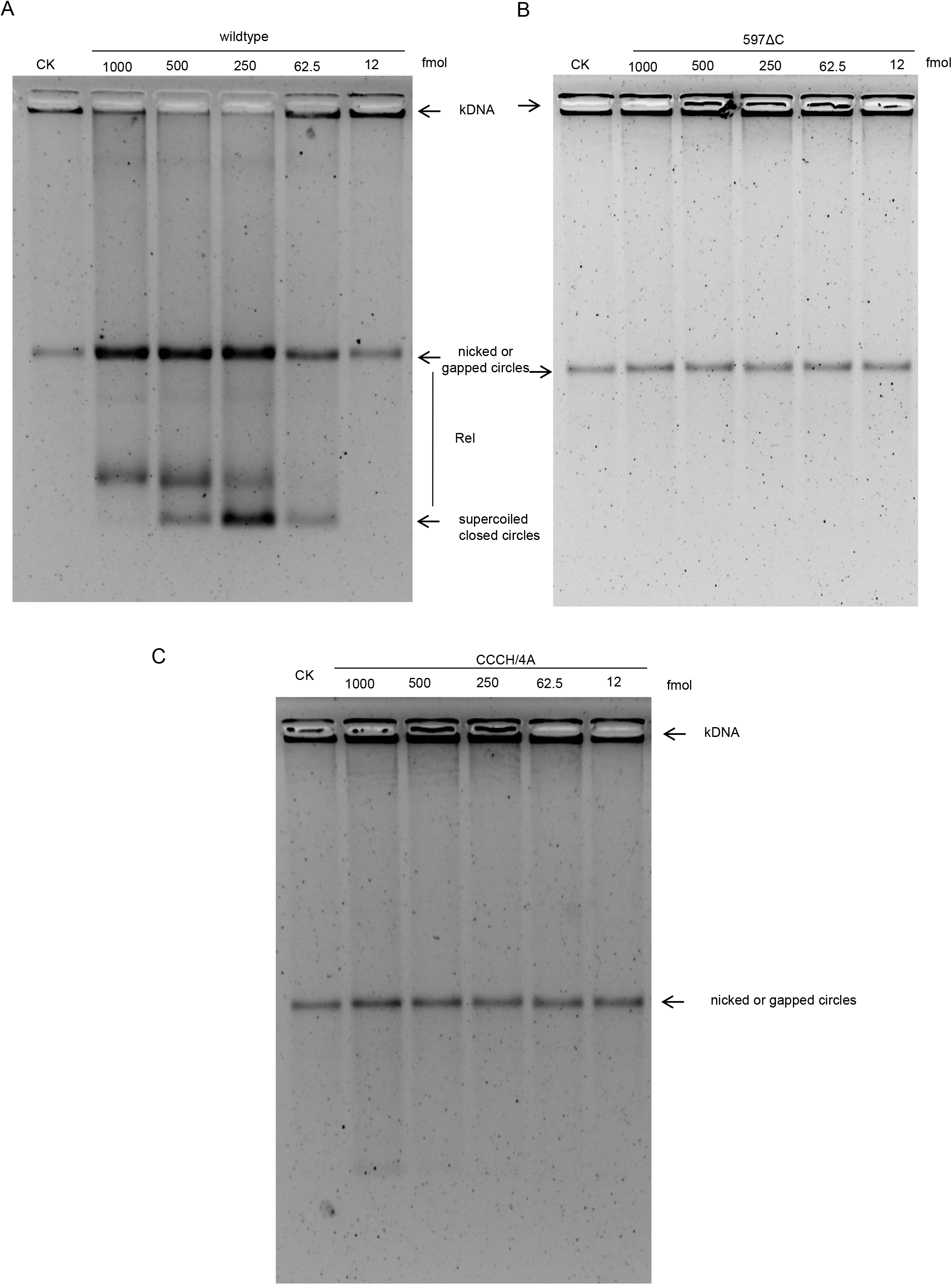
kinetoplast DNA (kDNA) decatenation by wild-type and mutants *Sso* topo III proteins. Decatenation assays were carried out as described for DNA relaxation assay except that kDNA (400 ng) was used as substrate with wild-type *Sso* topo III, 597ΔC and CCCH/4A mutant at 1000, 500, 250, 62.5 and 12 fmol. The reactions were performed at 90°C for 30 min and were terminated. Samples were electrophoresed through a 2% agarose gel, and the DNA products were visualized by staining with ethidium bromide. Lane CK corresponds to the kDNA control incubated at 90°C without the enzyme. Minicircles denote minicircle DNA released by *Sso* topo III.

## DISCUSSION

Type IA topoisoerases are found in all three domains of life. Much of current knowledge about the enzymes has come from structural and biochemical studies on the ones from bacterial and eukaryotic origins (41). A structural understanding of type IA topoisomerases (except for reverse gyrase) in the Archaea remains elusive. In this work, we present the crystal structure of *Sso* topo III, an archaeal type IA topoisomerase, at 2.1 Å resolution. The structure provides the first molecular view of this archaeal topoisomerase, which removes negative supercoils and decatenates DNA catenane (30, 32, 42). *Sso* topo III is toroidal in shape with a four-domain protein core closely resembling that found in other members of the topoisomerase IA family (6). A striking feature of *Sso* topo III is the presence at its carboxyl-terminus of a novel zinc finger-containing part that attaches to the hinge region of domain II. Intriguingly, the enzyme lacks the decatenation loop found in some of the known type IA topoisomerases capable of DNA decatenation. As revealed by mutational analysis, the zinc-binding motif is critical for effective decatenation of *Sso* topo III. Therefore, *Sso* topo III represents a novel type of topoisomerases which employ a DNA decatenation mechanism different from those of the known bacterial and eukaryotic topoisomerases III.

The carboxyl-terminal domain of a type IA topoisomerase varies widely in composition, length and shape (41, 43), as well as in orientations relative to the hinge region of the enzyme. Despite wide variations, the carboxyl-terminal domain exhibits ssDNA-binding activity, and is responsible for the high DNA-binding affinity of the enzyme. In the case of EcTOP1, domain IV extends into the carboxyl-terminal domain consisting of three zinc ribbon domains containing the tetracysteine motif and two zinc ribbon-like domains (39). Among the five domains, four domains bind to ssDNA with primarily π-stacking and cation-π interactions. Similarly, TmTOP1 has a zinc ribbon motif, but without a bound zinc ion, at its C-terminus (8). The deletion of the carboxyl-terminal domain resulted in a significant loss of the relaxation activity of TmTOP1, possibly due to the reduction in DNA-binding by the mutant protein (44, 45). By comparison, the truncation of the carboxyl-terminal domain almost abolished the relaxation activity of EcTOP3 (46), whose C-terminus displayed a partially disordered structure (7, 14). Unlike EcTOP1, MtTOP1 has four tandem novel folds, but not the zinc-binding motif, at the C-terminus (9). Similar to EcTOP3, the partial truncation of the carboxyl-terminal domain abolished the relaxation activity of MtTOP1. It is generally believed that the carboxyl-terminal domains of these two topoisomerases have ssDNA-binding activity, although they lack a zinc-binding motif. Our structure of *Sso* topo III provides the first molecular view of the carboxyl-terminal domain of an archaeal type IA topoisomerase (Fig. 1B). The carboxyl-terminal domain of *Sso* topo III is unique, and harbours a zinc-binding motif (3Cys1His type) (Fig. 1D) that differs from the tetracysteine zinc-binding motif conserved in prokaryotic type IA topoisomerases such as EcTOP1 (39). A structural analysis of the EcTOP1-DNA complex demonstrated that a zinc ribbon domain or zinc ribbon-like domain, mainly consisting of β-sheets, is capable of binding ssDNA, implying that these domains may not bind double-stranded DNA. In some enzymes involved in DNA transactions, the zinc-finger domains, which contain α-helixes, bind to double-stranded DNA (47). It will be interesting to determine whether the carboxyl-terminal fold of *Sso* topo III can bind double-stranded DNA. In *Sso* topo III, the zinc-binding motif is required for efficient relaxation activity (Fig. 3 and 4). By comparison, TmTOP1 (44) remained fully active in DNA relaxation when the zinc ribbon motif was deleted while EcTOP1 (48) was inactive when the zinc ion was removed.

The first solved structure of a type IA topoisomerase indicates that domains II and III may move away from domains I and IV in the strand passage reaction (6). Further evidence for this movement has been provided by biochemical (19, 21), structural data (14, 15, 35) and single-molecule experiments (22–24). A structural comparison between EcTOP3 and EcTOP1 indicates that there are two unique insertions in EcTOP3 in the vicinity of its central hole (29) (Fig. 2A). One insertion (the decatenation loop) is rich in basic amino acids within domain IV of EcTOP3, and the other (the acidic loop) is abundant in acidic amino acids and is located within domain II (2) (Fig. 2A). Decatenation by the enzyme is driven primarily by the interaction between the decatenation loop and the acidic loop, both of which line the central hole of the topoisomerase, during its gate dynamics. Compared with the interaction in EcTOP1, that in EcTOP3 presumably stabilizes and holds the open state for longer, thus allowing sufficient time for EcTOP3 to catch duplex DNA and pass it through the break in a single strand into the gate (24). In the eukaryotic Top3a-RMI complex, RMI1 coordinates the gate dynamics of Top3a through its unique decatenation loop, which has a functionally equivalent role to that of the decatenation loop in EcTOP3 (11). Hence, a unified loop-mediated decatenation mechanism for prokaryotic and eukaryotic topoisomerases III has been proposed (11).

Our results demonstrated that the zinc-binding motif is essential for the DNA decatenation activity of *Sso* topo III (Fig. 5C). We provided the following evidence in support of the notion that the decatenation mechanism of *Sso* topo III is distinct from the current loop-mediated decatenation mechanism proposed for type IA topoisomerase: (i) The decatenation domain (domain V) of *Sso* topo III is a novel zinc finger fold (Fig. 1D), whereas the decatenation loop is present in domain IV of EcTOP3 (29) (Fig. 2A). (ii) The decatenation domain (domain V) of *Sso* topo III is on the outer edge of the topoisomerase gate and attaches to domain II through the hinge region by extensive hydrophobic interactions (Fig. 1D and 2C), whereas the decatenation loop in EcTOP3 lines the DNA gate and is away from the hinge region of the catalytic core (24) (Fig. 2A). (iii) The 594ΔC mutant lacking domain V showed an approximately 5-fold reduction in ssDNA-binding compared with wild-type *Sso* topo III (Fig. S5). In contrast, the deletion of the decatenation loop from EcTOP3 did not reduce its ssDNA-binding activity (29). Obviously, further biochemical studies are needed to understand the molecular details of the unlinking of covalently closed catenanes catalysed by *Sso* topo III.

## MATERIALS AND METHODS

### Plasmid and DNA constructs

Using the standard PCR cloning strategy, *Sso* topo III gene from *S. solfataricus* genome was cloned into the *NdeI* and *XhoI* sites of the modified version of vector pET28a (Novagene), which harbored an N-terminal poly-histidine tag followed by a tobacco etch virus (TEV) protease cleavage site (ENLYFQ/G) (where ‘/’ donates the cleavage site). However, the growth of *E.coli* cells harboring wild-type *Sso* topo III gene were severely inhibited, possibly due to its toxicity. So, *Sso* topo III gene was mutated to the *Sso* topo III Y318F by replacing residue Tyr318 with Phe using the QuickChange^™^ Site-Directed Mutagenesis Kit (Stratagene) according to the instructions of manufacturer. Site-directed mutagenesis of the other *Sso* Topo III mutants used in this study was carried out according to modified QuickChange^™^ protocol (49). All the plasmid constructs and mutants were confirmed by DNA double strands sequencing. Primers for *Sso* Topo III site-directed mutagenesis are listed in Table S1.

### Protein Expression and Purification

The recombination *Sso* topo III Y318F was overexpressed as a fusion protein with an N-terminal poly-histidine tag in *E. coli* strain Rosetta2 (DE3) pLysS (Novagen). For each purification procedure, 10-l *E. coli* cultures harbouring fusion proteins were grown in the shaker at 37°C at 185 r.p.m. When their density reached an OD_600nm_ of 1.2, sodium citrate was added into cultures to a final concentration of 100 mM and temperature of the cultures dropped to 25°C, followed by the addition of isopropylb-D-thiogalactoside to a final concentration of 0.5 mM.

After growth for further 6 h at 25°C, the cultures were harvested, and the resulting cell sediments were suspended in a solubilization buffer containing 20 mM Tris-HCl (pH 8.0), 2 mM potassium citrate, 2 mM MgCl2, 10 mM imidazole and 500 mM NaCl supplemented with DNAase I (55 μg/ml, Amersco), followed by disruption with a French press at 12,000 p.s.i. for three cycles. To remove cell debris, cell lysates were centrifuged for 40 min at 16,000 r.p.m. The resultant supernatant was collected and loaded onto an equilibrated Ni^2+^ Chelating Sepharose^™^ Fast Flow column (GE Healthcare). Purification procedure with the nickel column were performed as described before (50), except the insertion of an additional wash with high-salt of 1 M NaCl. After several wash, *Sso* topo III Y318F protein was eluted with a buffer containing 20 mM Tris-HCl (pH 8.0), 2 mM potassium citrate, and 300 mM NaCl supplemented with 300 mM imidazole. The eluate was digested using TEV protease to remove the poly-histidine tag, and dialyzed against buffer I (20 mM Tris-HCl, pH 7.5, 10 mM potassium citrate, 150 mM NaCl) overnight at 4°C. The resultant eluate was filtered through a 0.45 μm filtration membrane and then loaded onto a 5 ml SOURCE 15S column (GE Healthcare) equilibrated with buffer I. The column was eluted with an 80 ml linear gradient from 150 mM NaCl to 500 mM NaCl, both in buffer I. The fractions, eluted at 0.45-0.50 M NaCl, were pooled and concentrated to 4 mg ml^−1^ in a buffer containing 20 mM Tris-HCl (pH7.5), 150 mM NaCl, 10 mM potassium citrate, and 2 mM DTT for later usage. Purified protein was analysed by SDS-PAGE for purity and quantified with Bradford assay using BSA as standards.

Expression and purification of the other *Sso* Topo III mutants (597ΔC and CCCH/4A) were carried out according to the procedure described above, with the exception of ion-exchange column chromatography in the final purification step. The resultant nickel column eluate was loaded onto a 5 ml SOURCE 15Q column (GE Healthcare), instead of 15S column, equilibrated with buffer I. The column was eluted with an 80 ml linear gradient from 0.150 mM NaCl to 1.0 M NaCl, both in buffer I. The fractions, eluted at 0.55-0.65 M NaCl, were pooled.

### Crystallization

*Sso* topo III Y318F was crystallized by the sitting-drop vapor diffusion method, with 0.7μl drops of protein or protein-ssDNA complex mixed with an equal volume of reservoir solution. Initial crystallization trials screen clustered-needle-shaped crystals in reservoir solution containing 8% (v/v) Tacsimate (pH 4.4), and 20% (w/v) Polyethyleneglycerol 3,350. Further optimization of the crystallization condition led to the appearance of rod-shaped crystals in the well in which 0.7μl protein solution was mixed with an equal volume of reservoir solution consisting of 8% (v/v) Tacsimate (pH 4.4), and 13.75% (w/v) Polyethyleneglycerol 3,350 at 12°C. The diffraction-level crystals, by the microseeding method, appeared one week later and reached a maximum size within 2 months. These crystals in the mother liquor were exposed to air for 10 min, and transferred into a drop of paraffin and NVH mix oil before being crycooled by plunging rapidly in liquid nitrogen.

### Crystallographic Data Collection, Structure determination and Refinement

X-ray diffraction data were collected at the BL17U beamline of the Shanghai Synchrotron Radiation Facility, and indexed, integrated and scaled using the HKL2000 package (51). The initial phases were determined by molecular replacement using the program PHASER (52), with the structure of TmTOP1 (PDB ID: 2GAJ) as a searching model. The structure refinement was carried out with phenix.refine (53) and Refmac5 (54). Model building was carried out by iterative rounds of manual building with Coot (55). MolProbity (56) was used to validate the structure. Data collection and refinement statistics are listed in Table 1.

### Assays of decatenation and relaxation activities

DNA relaxation assay was performed as described previously (30), except that the final concentration of MgCl2 is 20 mM, instead of 10 mM, in the standard reaction mixture. Decatenation of kinetoplast DNA (kDNA) was carried out by a similar protocol except that 400 ng kDNA (www.topogen.com) was used as substrate in each standard reaction mixture. Reaction mixes were incubated at 90 °C for 30 min unless specified temperature otherwise, and the enzyme-catalysed products were examined by electrophoresis in the indicated agarose gels.

### Data availability

Atomic coordinates and structure factors for the crystal structure of *Sso* topo (apo) have been deposited in the Protein Data bank under accession codes 6K8N.

## SUPPORTING INFORMATION

Additional supporting information may be found online in the Supporting Information section.

## ACKNOWLEDGEMENTS

We are grateful to Dr. Mei Li and the staff of beamline BL17U at the Shanghai Synchrotron Radiation Facility and of beamline BL5A at the Photon Factory, KEK (Tsukuba, Japan) for technical support. This study was financially supported by National Natural Science Foundation of China [U1432241 to Y.G. and Z.Z., 31730001 and 91751000 to L.H., 92051109 to Z.Z.]; Strategic Priority Research Program of the Chinese Academy of Sciences [XDB37040302 to Y.D.]; National Basic Research Program of China [2017YFA0504900 to Y.D.]; and Beijing Municipal Science and Technology Commission [ Z191100007219007 to Y.D.]. Funding for open access charges was provided by the National Natural Science Foundation of China.

We declare no conflict of interest.

