## Supplemental Table 1 for "Crystal structure of *Sulfolobus solfataricus* topoisomerase III reveals a novel carboxyl-terminal zinc finger domain essential for decatenation activity"

Supplementary Information

Table S1 Primers for SsTopIII site-directed mutagenesis

| Mutagenesis | Primer name | Primer sequence^a^ (5' to 3') |
| --- | --- | --- |
| Y318F | Y318F-F | *GGACGGTCTAATAAGT***TTC***CCAAGAACTAACAGTC* |
|  | Y318F-R | *GACTGTTAGTTCTTGG***GAA***ACTTATTAGACCGTCC* |
| 597ΔC (I597dele) | I597dele-F | *AGGAGAGTCATTAGCTAAA*GCGTTAGGTCTTATA**TAATAA**TTGTTAAGTG |
|  | I597dele-R | *TTTAGCTAATGACTCTCCT*ACTTTATCCTTATTTACCTTGTATTCCTC |
| CCCH/4A (C602S/C605S/C615S/H618A) | C602S/C605S/C615S/H618A-F | *GGAGCAGTATAAGGATG*GATTA**TCT**AAATAT**GCC**TATGAAGCCAAAGTAAGG |
|  | C602S/C605S/C615S/H618A-R | *CATCCTTATACTGCTCC*AGATC**AGA**ATACTT**AGA**CTTAACAATTTTTATAAGACC |

a. Sense primer-antisense primer overlapping sequences are in italic. Codons encoding mutated residues and stop codon are in bold.

Supplementary Figure legends

FIG S1. Topology cartoon presentation of the structure of *Sso* topoIII.

Domains are color-coded: domain I (purple), II (green). III (blue), IV (yellow) and V (red). Cysteine residues 4, 34, 602, 605, and 615, and histidine residue 618 are shown with cycles. Cysteine residues 602, 605, 615 and histidine residue 618 coordinate zinc atom in domain V, and cysteine residue 4 links 34 to form disulfide bond in domain I.

FIG S2. Structure-based amino-acid sequence alignment of regions, mediating interface between domain V and domain II, among topoisomerase from archae.

The secondary structure elements of *S. so* topoisomerase are shown above the alignment. Arrows indicate β-strands and coils indicate α helices.

Strictly conserved residues are boxed and marked with red back ground and conservatively substituted residues are boxed. *Sso*Topo3，*Sulfolobus solfataricus* Topoisomerase III; SisTopo3, *Sulfolobus islandicus* Topoisomerase III; StoTopo3, *Sulfolobus tokodaii* Topoisomerase III; AhoTopo1, *Acidianus hospitalis* W1 Topoisomerase I; AhoTopo3, *Acidianus hospitalis* Topoisomerase III; SacTopo1, *Sulfolobus acidocaldarius* Topoisomerase I; SacTopo3, *Sulfolobus acidocaldarius* Topoisomerase III;

CacTopo1, *Candidatus Acidianus copahuensis* Topoisomerase I; SarTopo1, *Sulfolobales archaeon* AZ1 Topoisomerase I; TagTopo1, *Thermosphaera aggregans* Topoisomerase I; SheTopo1, *Staphylothermus hellenicus* Topoisomerase I; TceTopo1, *Thermogladius cellulolyticus* Topoisomerase I;

SmaTopo1, *Staphylothermus marinus* Topoisomerase I; PdeTopo3, *Pyrodictium delaneyi* Topoisomerase III; HbuTopo3, *Hyperthermus butylicus* Topoisomerase III; DmuTopo1, *Desulfurococcus mucosus* DSM 2162 Topoisomerase I; IhoTopo1, *Ignicoccus hospitalis* Topoisomerase I;

IagTopo1, *Ignisphaera aggregans* DSM 17230 Topoisomerase I;

A. The conserved amino acid residues in zinc finger motif [C3H1] are indicated with red “*”; In domain V, the residues forming hydrogen bonds and hydrophobic interaction with those from domain II are shown with red dots and blocks below the alignment, respectively.

B. In domain II, the residues forming hydrogen bonds and hydrophobic interaction with those from domain V are marked with black dots and blocks below the alignment, respectively.

FIG S3. Phylogenetic profile of domain V of topoisomerase III from *Sulfolobus solfataricus* in Archaea.

Homologues of *Sso* topo III from Crenarchaeota are shown in blue, and domain V-like short proteins from Euryarchaeota, Thorarchaeota, Bathyarchaeota and Thaumarchaeota are shown in red. The total number of Domain V homologues identified in each taxon is indicated.

FIG S4. Effect of temperature on DNA relaxation by wild-type and mutant *Sso* topo III proteins.

DNA relaxation assays were performed at the incremental temperatures as described in the legend to FIG 3. The individual assay temperature is indicated above each panel. The protein/ pUC18 DNA molar ratio used was 1:1. The wild-type (A), 597ΔC (B) and CCCH/4A (C) were subjected to DNA relaxation assay. The reaction products were analyzed in 1.4% agarose gels. OC, open circle (nicked or gapped circular) DNA, Rel, the relaxed topoisomers, and SC, negatively super-coiled circular DNA. FI* corresponds to unlinked DNA species, and were apparently identical to highly unwound species.

FIG S5. DNA binding by the carboxyl-terminally truncated mutant 594ΔC of *Sso* topo III.

Wild-type *Sso* topo III or 594ΔC (1.0, 5.2, 26.1 and 131 nM) was incubated at 25 °C for 5 min with ^32^P-labeled C32 (1.25 nM) in the absence of MgCl_2_. Samples were subjected to electrophoresis in 8% polyacrylamide gel. The gel was exposed to X-ray film. Oligonucleotide C32, 5′-GCCCTTGGCAAGGTCTCCCCCCCCTTTTTTAT-3’) (the major recognition sequence for cleavage is underlined) was employed as the binding strand for *Sso* topo III. C32 was labeled at the 5′ end with [ϒ-^32^P] ATP using T4 polynucleotide kinase.

FIG S6. Effect of temperature on DNA decatenation by wild-type *Sso* topo III.

A. DNA decatenation assays were performed at the incremental temperatures as described in the legend to FIG 5. 250 fmol of protein was used in the assay. The individual assay temperature is indicated above each panel. The reaction products were analyzed in 1.3% agarose gel.

B. DNA decatenation by wild-type *Sso* topo III at 85°C. DNA decatenation assays were performed as described in the legend to FIG 5. The reaction products were analyzed in 2% agarose gel.

C. DNA decatenation by wild-type *Sso* topo III at 95°C. DNA decatenation assays were performed as described in the legend to FIG 5. The reaction products were analyzed in 1.3% agarose gel.
