## Supplementary figures and images for "Crystal structure of *Sulfolobus solfataricus* topoisomerase III reveals a novel carboxyl-terminal zinc finger domain essential for decatenation activity"

### Supplemental Fig. 1

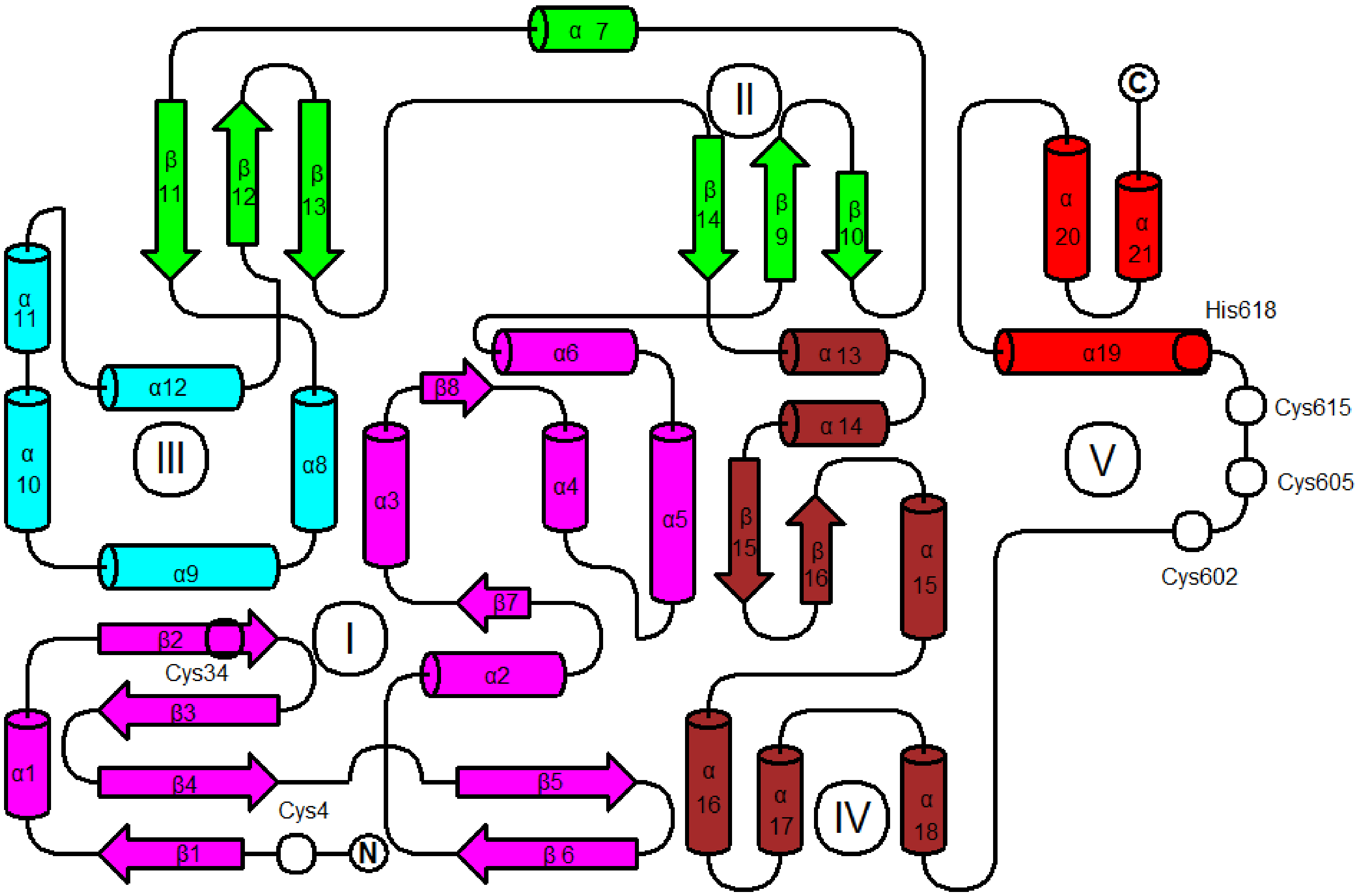

### Supplemental Fig. 2

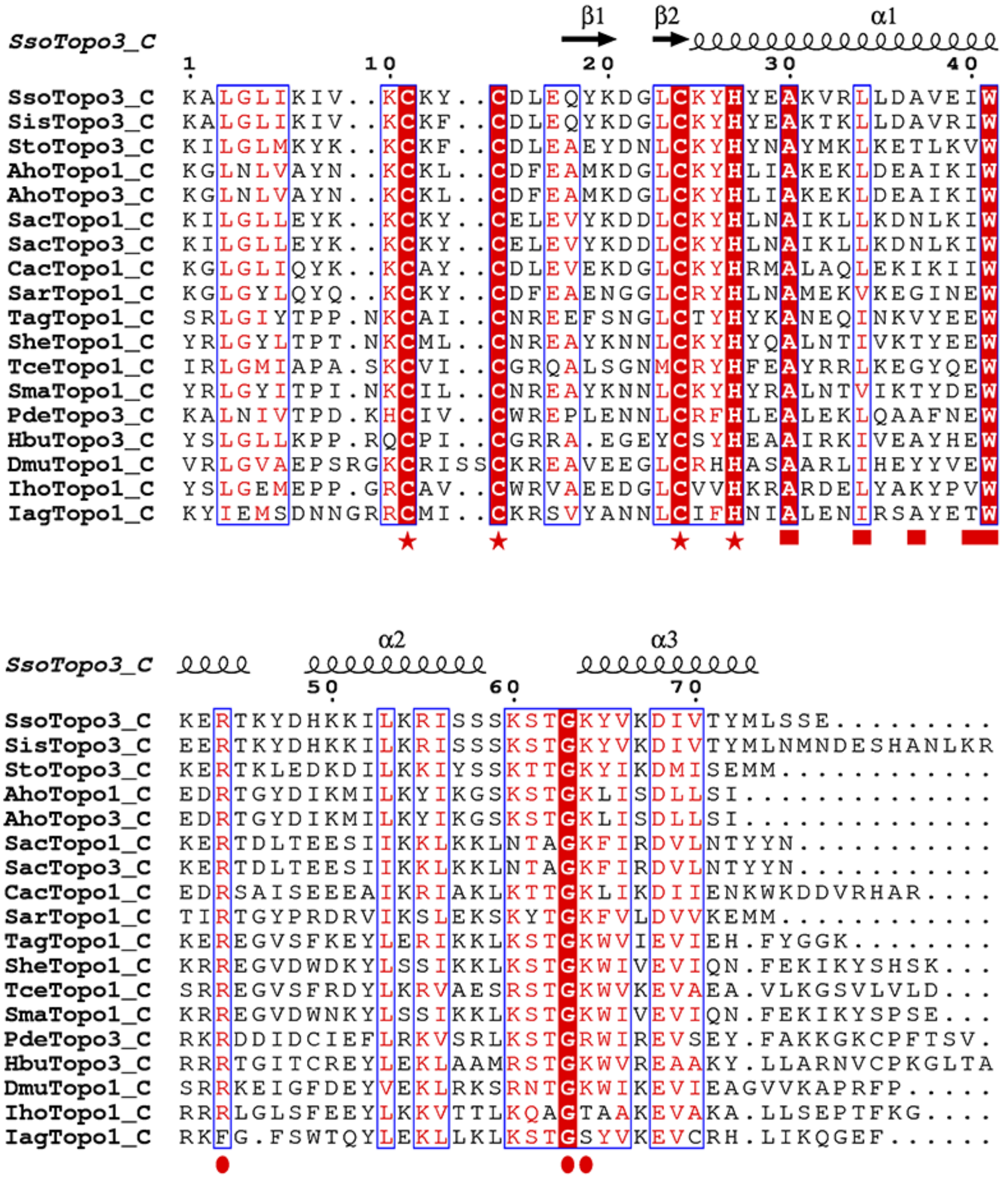

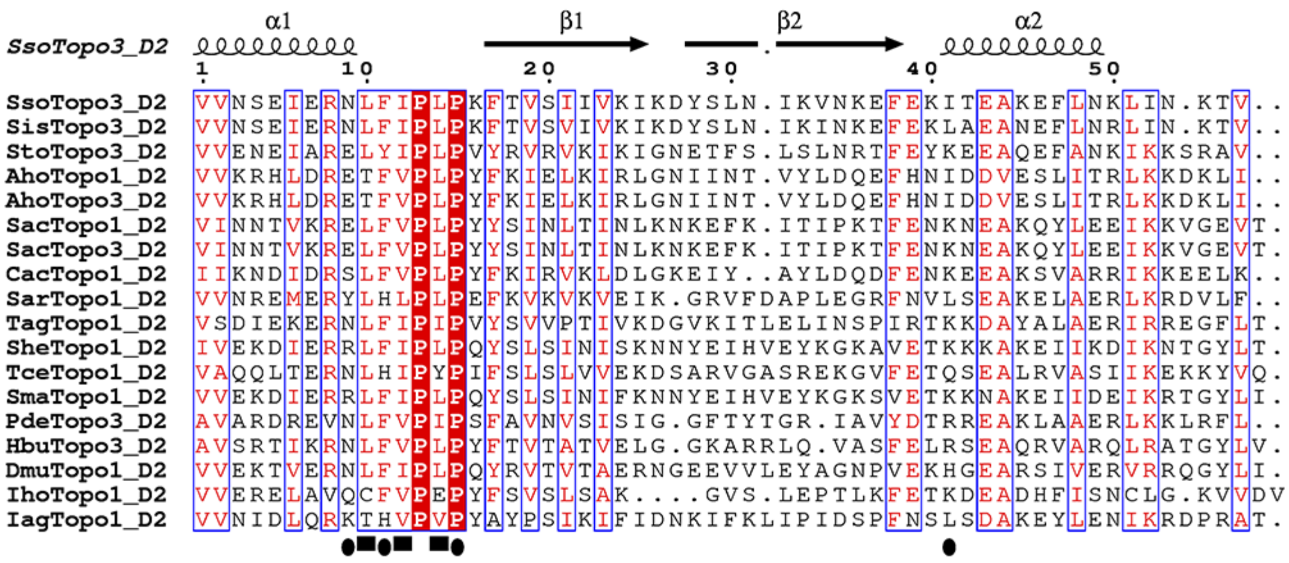

### Supplemental Fig. 3

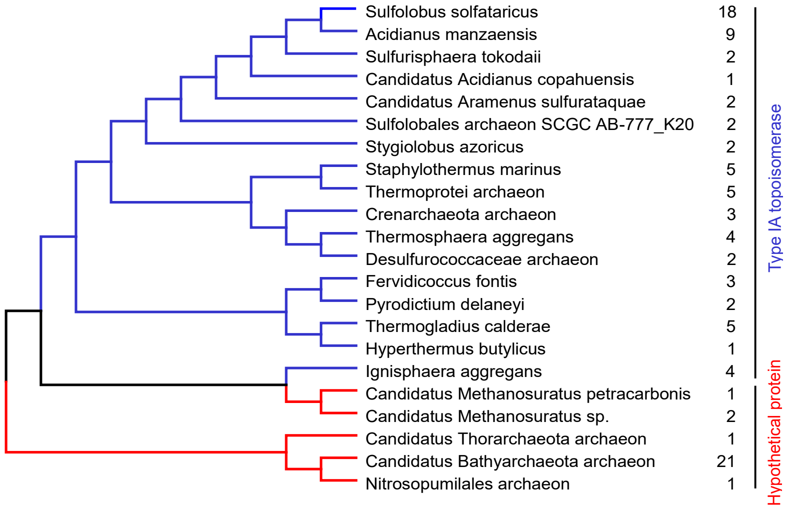

### Supplemental Fig. 4

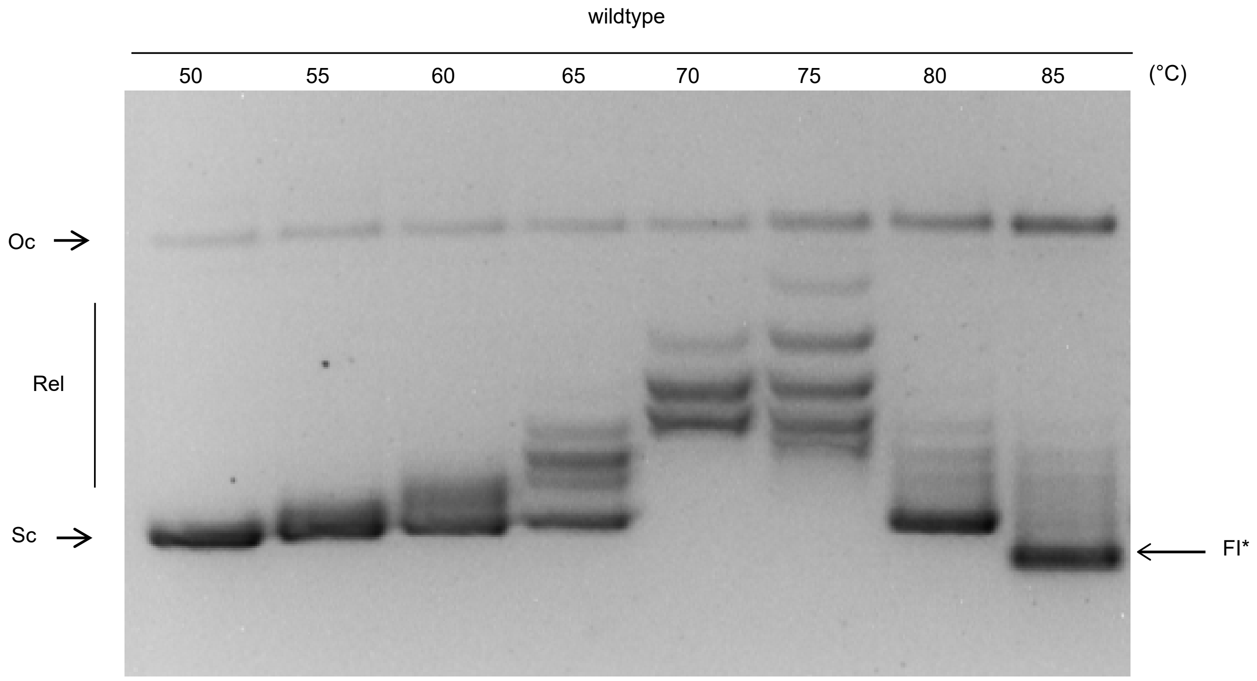

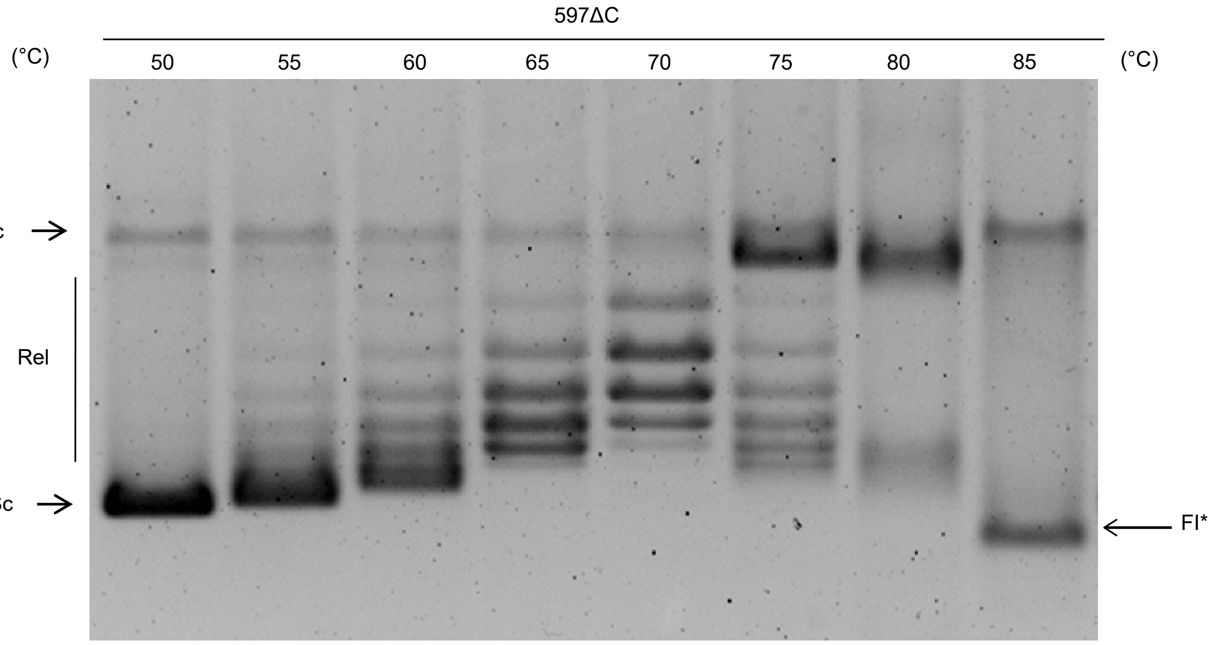

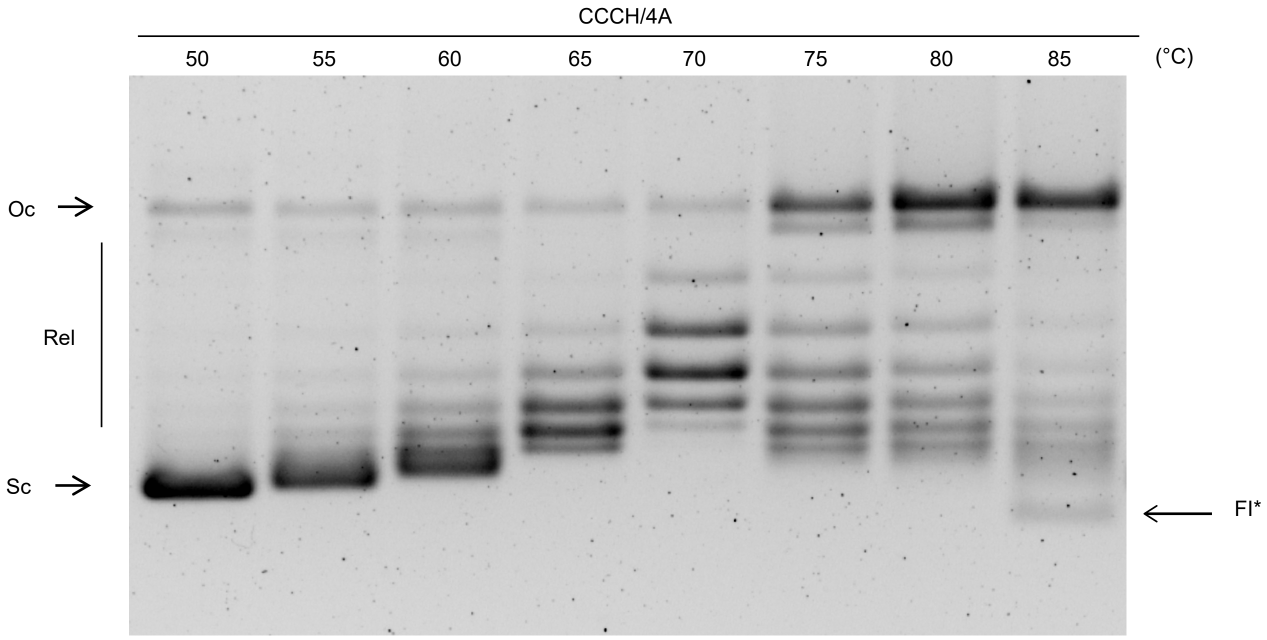

### Supplemental Fig. 5

FIG S5

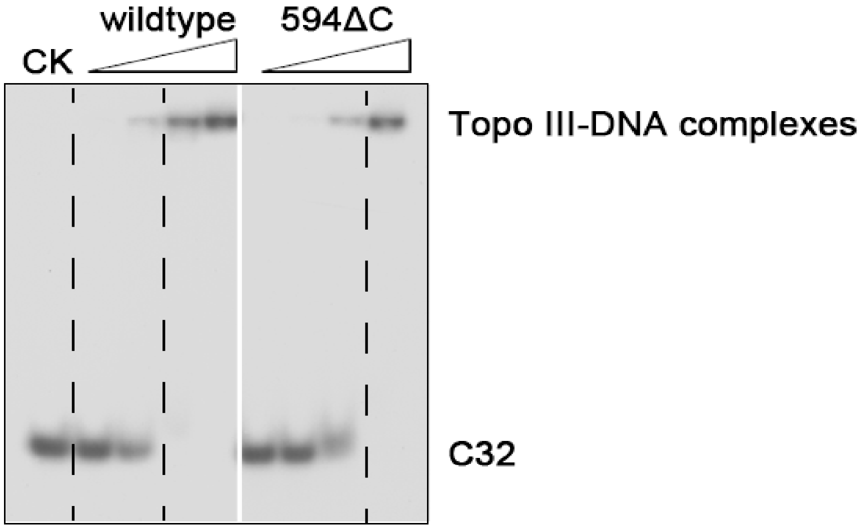

### Supplemental Fig. 6

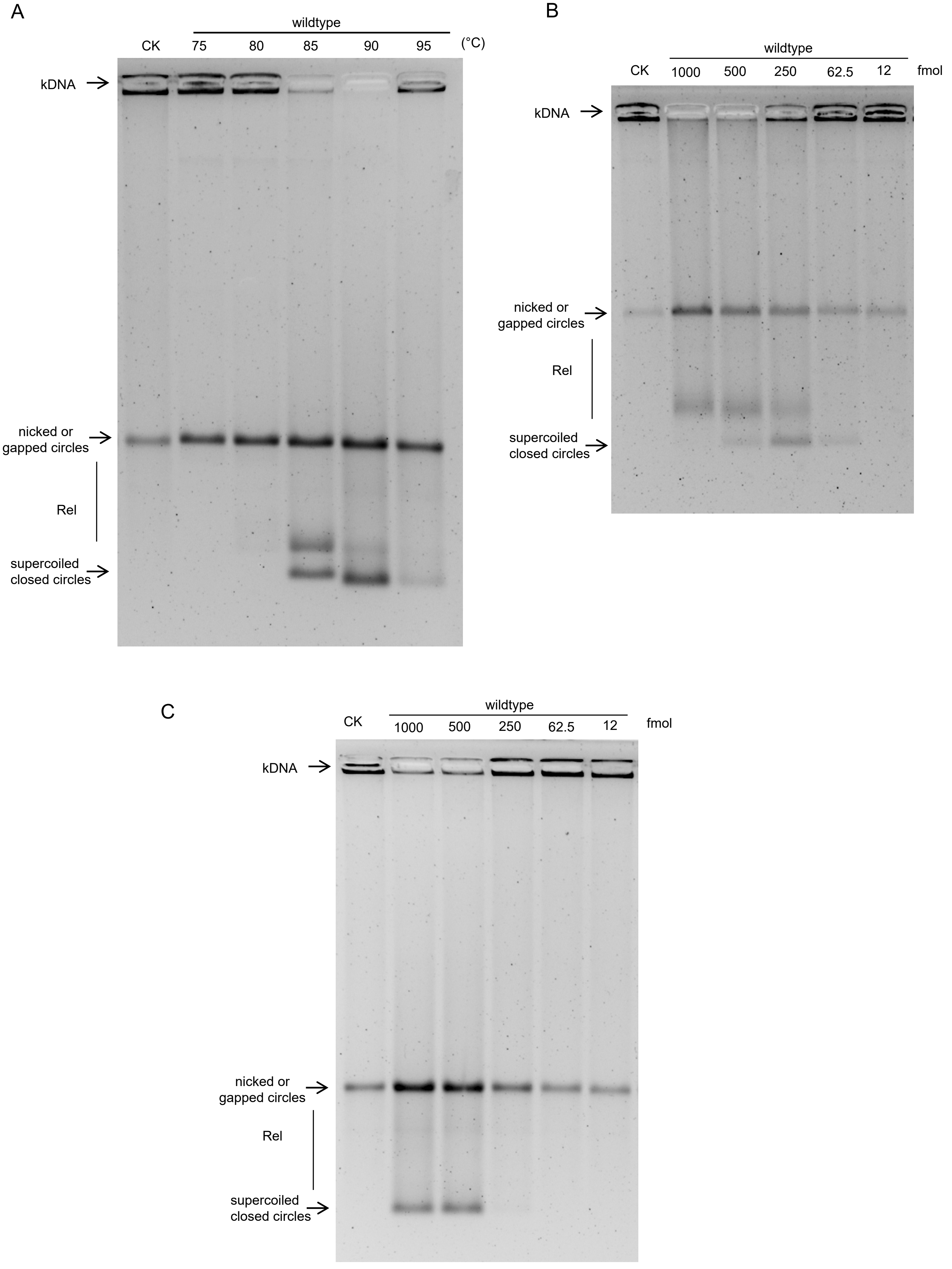
